# Molecular dynamics simulation study reveals effects of key mutations on spike protein structure in SARS-CoV-2

**DOI:** 10.1101/2021.02.03.429495

**Authors:** Jerome Rumdon Lon, Binbin Xi, Bingxu Zhong, Yiyuan Zheng, Zixi Chen, Ruoran Qiu, Siqing Zhang, Pei Guo, Hongli Du

**Affiliations:** School of Biology and Biological Engineering, South China University of Technology, Guangzhou, China; Institute of Synthetic Biology, Shenzhen Institute of Advanced Technology, Shenzhen, China

**Keywords:** Virology, Molecular biology, Molecular dynamics, Bioinformatics

## Abstract

SARS-CoV-2 has been spreading rapidly since 2019 and has produced large-scale mutations in the genomes. The mutation in genes may lead to changes in protein structure, which would have a great impact on the epidemiological characteristics. In this study, we selected the key mutations of SARS-CoV-2 from a real-time monitoring tool, including D614G, A222V, N501Y, T716I, S982A, D1118H of spike (S) protein, and performed molecular dynamics (MD) simulations on single-site mutant D614G, double-site mutant D614G&A222V and penta-site mutant N501Y&D614G&T716I&S982A&D1118H to investigate their effects on protein structure and stability using molecular dynamics (MD) simulations. The results suggested that D614G improved the stability of S protein, while D614G&A222V and N501Y&D614G&T716I&S982A&D1118H showed an increased solvent accessible surface area and they might enhance the ability of protein to react with the outside environment. Our findings could complement the mechanistic link between genotype--phenotype--epidemiological characteristics in the study of SARS-CoV-2. We also found no significant difference between the antigenicity of S protein and the mutants through Ellipro, which may reference for vaccine development and application.

## Introduction

In the winter of 2019, a new coronavirus, SARS-Cov-2, came into view and quickly sprea<u>d</u> the world. By 0:00 on June 16th, 2021 (GMT+8:00), the total number of diagnosed patients has exceeded 176.6 million. SARS-Cov-2 has been a major blow to public health, social order, and economic development around the world.

As a transmembrane protein, the S protein mediates coronavirus to entry into host cells. Owing to its important role in virus transmission ^1^, S protein has been the focus of coronavirus-related research studies. The S protein forms a homotrimer on the surface of the virus, containing the S1 subunit that binds to the host cell receptor and the S2 subunit that is involved in the fusion of the viral membrane to the cell membrane. A series of studies have shown that two subunits of the S protein of coronavirus bind non-covalently in a prefusion conformation, while host proteases lyse the S protein during viral entry into the cell. This modification has been proposed to activate the protein for membrane fusion by irreversible conformational changes ^2–5^. Because the S protein is exposed to the surface and plays an important role in transmission, it is considered to be the primary target of post-infection neutralizing antibodies and is the focus of treatment and vaccine design. Due to its important role in viral mechanism and prevention measures, the mutation in the S protein is likely to have an impact on the phenotype and epidemiological characteristics of the virus, and perhaps the function of antibodies.

As the virus spreads, it accumulates a large number of genomic mutations that could alter the structure and function of proteins, which would increase the complexity of the virus subpopulation and pose a threat to prevention and control measures such as virus detection, drug development, and the practical application of vaccines. In the case of MERS-CoV, the I529T, D510G mutations in RBD lead to decreased affinity with the human CD26 receptor ^6^, but increased resistance to antibody-mediated neutralization. In the case of SARS-CoV ^7^, the P462L mutation of S protein can achieve the escape of human mAb Cr3014 ^8^. Recently, the United Kingdom, South Africa, Nigeria, and other places have reported the emergence and prevalence of mutant strains ^9–12^. More disturbingly, a SARS-CoV-2 variant that achieved *in vitro* escapement from highly neutralizing COVID-19 convalescent plasma has been reported ^13^. Therefore, a real-time monitoring of the epidemic trends of key mutations and haplotypes in virus evolution, and exploring the influence of mutations on virus phenotypes, as well as the relationship between mutations and epidemiological characteristics are of great significance for the development of targeted prevention and control measures. Our previous research has revealed six key mutations in the spike(S) protein, including the D614G, A222V, N501Y, T716I, S982A, and D1118H^14–16^. The single D614G mutation variant began to spread in Europe in early February of 2020, which has been reported to significantly increase the infectivity of SARS-CoV-2 in continuous cell lines ^17–21^. The D614G&A222V mutation variant was firstly discovered from Spain in March of 2020 and then rapidly spread since July across Europe and accounted for about 16.3% of all GISAID submitted genomes ^15^. The other four mutations in S protein, including N501Y, T716I, S982A, and D1118H, were also detected from the subsequent monitoring of SARS-CoV-2 transmission by our real-time monitoring tool^16^, which had also been mentioned previously^9^. The four mutations were almost completely linked and mutated based on the single D614G mutation variant^16^, These imply the D614G&A222V variant and the N501Y&D614G&T716I&S982A&D1118H variant may also play an important role in increasing infectivity of SARS-CoV-2.

Based on the real-time monitoring of pandemic virus mutants^15,16^, this study conducted molecular dynamics (MD) simulations of S protein based on the linkage mutations rather than single by single mutation sites, which could better simulate the real situation of pandemic mutants. Then, we have performed MD simulations to evaluate the effect of D614G single-site mutation, D614G&A222V double-site mutation, and the N501Y&D614G&T716I&S982A&D1118H penta-site mutation on the structural properties of S protein, as A222V and N501Y&T716I&S982A&D1118H have been reported to link with D614G respectively ^15,16^. The results showed that, comparing with the wile type (WT) S protein, the RMSD values of D614G, D614G&A222V, N501Y&-D614G&T716I&S982A&D1118H were significantly decreased and their corresponding stability was increased. However, the radius gyration (R_g_) and solvent accessible surface area (SASA) of D614G and D614G&A222V were found to be lower than those of WT, while the R_g_ and SASA of penta-site mutation appeared to be similar to the WT. In addition, we also analyzed the antigenicity of D614G, D614G&A222V and N501Y&D614G&-T716I&S982A&D1118H, it was found that the structural change caused by site mutations had no significant effect on the antigenicity of these S protein mutants. Yet, but the potential consequences of local changes, such as reinfection, are still worth attention. Taken together, our results provide new insights into the relationship of “genotype-phenotypic-epidemiological characteristics” of SARS-CoV-2. In particular, this study simulates multiple mutation sites coupling in the current virus transmission process, which will help to better understand the virus and address the complexities of COVID-19 prevention, diagnosis, and treatment.

## Results

### Analysis of six mutations and their potential influences on S protein structure and function

For the 6 site mutations of S protein studied in this work including D614G, A222V, N501Y, T716I, S982A, D1118H (Figure 1, Supplementary Information Movie 1). Expect T716I, the others all locate in the interface or boundaries between two chains or subunits, which are crucial positions for maintaining the structure and stability of the S protein structure. Therefore, we speculated that these mutations may affect the structure and morphology of S protein. Through a more detailed analysis on the high-resolution structure of the S protein (PDB ID: 6×6P)^22^, D614G and T716I are close to the junction site between S1 and S2 subunits ^23,24^, which may affect the processing and modification of S protein through the change of steric hindrance. A222V is located at the edge of the head of S protein and is close to the Receptor binding domain (RBD), while N501Y is located at the receptor-binding motif of RBD, which may also affect the protein properties. S982A is also located at the binding site between chains, and the loss of hydroxyl in the side chain caused by this mutation may have an effect on the interaction between chains. D1118H is located between two heptad repeats, and changes in the side chain may affect the interaction between the chains through spatial effects, thus affecting the properties of the whole protein.

**Figure 1.**
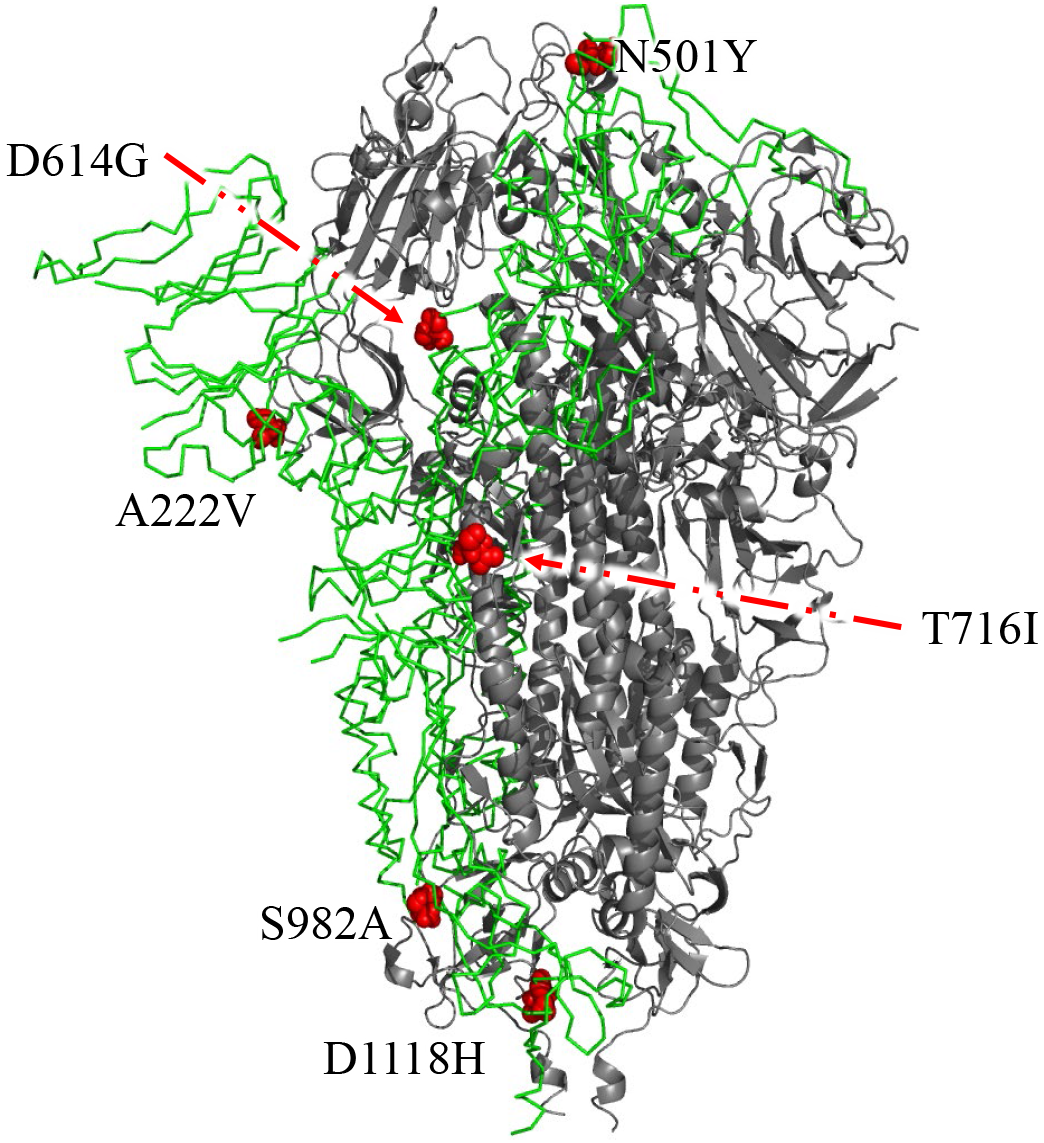
The structure of S protein(trimer) and the position of mutation sites.

### The stability of single, double, and penta-site mutants

Based on the reported cryo-electron microscopy structure of S protein (PDB ID: 6×6P)^22^, we constructed single-site D614G, double-site D614G&A222V, and penta-site N501Y&D614G&T716I&S982A&D1118H mutants. We exploited 100-ns MD simulations for the D614G, D614G&A222V, N501Y&D614G&T716I&S982A-&D1118H, and wide type (WT) S proteins. To verify the convergence of MD simulation equilibrium, we first analyzed the root mean square deviations (RMSD) throughout the MD process (Figure 2A), after excluding the initial effect of model relaxation due to the introduction of the force field in the first half of simulation^25^. The RMSD of the WT, D614G, D614G&A222V, and N501Y&D614G&T716I&S982A&D1118H mutants appeared to be stable in the last 50ns, revealing a good convergence for each system. In the last 50 ns which can be considered as a stable period (Table S1), the RMSD of D614G, D614G&A222V, and N501Y&D614G&T716I&S982A&D1118H appeared to be a bit smaller than that of the WT, and the difference became even more obvious in the last 25 ns. For the three mutants, their RMSD fluctuated within a small range and did not show an obvious difference in the last 25 ns.

**Figure 2.**
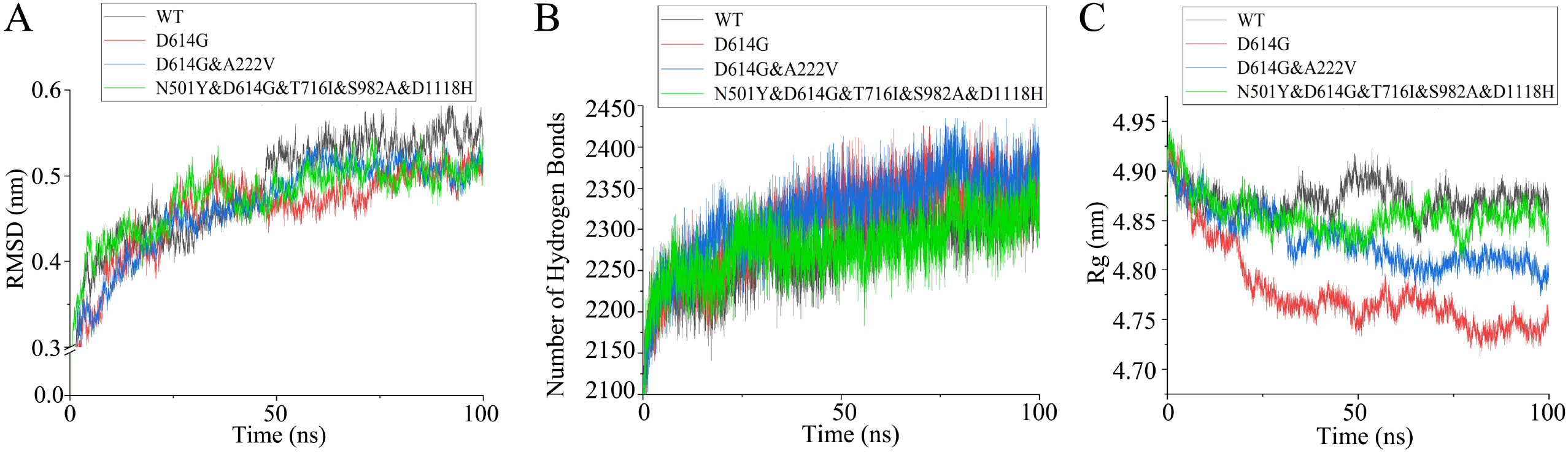
Stability evaluation of WT and its mutants of S protein. A. The RMSD of S protein and mutants. B. The H-bond of S protein and mutants. C. The R_g_ of S protein and mutantsFigure 3 Evaluation of the flexibility of WT and its mutants of S protein

We also performed the hydrogen bond analysis for the three mutants and WT (Table S2). The average number of hydrogen bonds follow the order of D614G&A222V > D614G > WT > N501Y&D614G&T716I&S982A&D1118H (Figure 2B). These results suggest that the D614G and D614G&A222V mutants are likely to be more stable than the WT. The structural compactness of the WT and mutants was studied by evaluating the R_g_ which provides information for the overall size of the structure (Table S3). The Rg values follow the order of WT ≈ N501Y&D614G-&T716I&S982A&D1118H > D614G&222V > D614G (Figure 2C), suggesting that the structure of D614G is more compact than other mutants and the WT.

To further investigate the structural dynamics of the D614G, D614G&A222V, and N501Y&D614G&T716I&S982A&D1118H mutants, we evaluated the residual atomic fluctuations by computing the root mean square fluctuations (RMSF) of protein Cα atoms (Figure 3, Table S4). It should be noted that the RMSF of highly dynamic terminal residues is not shown here. RMSF analysis showed that the overall trend of the flexibility of the WT and three mutants did not change too much. The RMSF at position 614 has a little difference between WT, D614G (Figure 3A), and other mutants (Figure 3B-3C). The D614G&A222V mutant showed a steep increase in local structure flexibility at the 222^th^ amino acid position (Figure 3B). Interestingly, all of the three mutants showed a steep reduced RMSF at the 501^th^ amino acid position, which may correlate with D614G as it appeared in the three mutants(Figure 3A-C).

**Figure 3.**
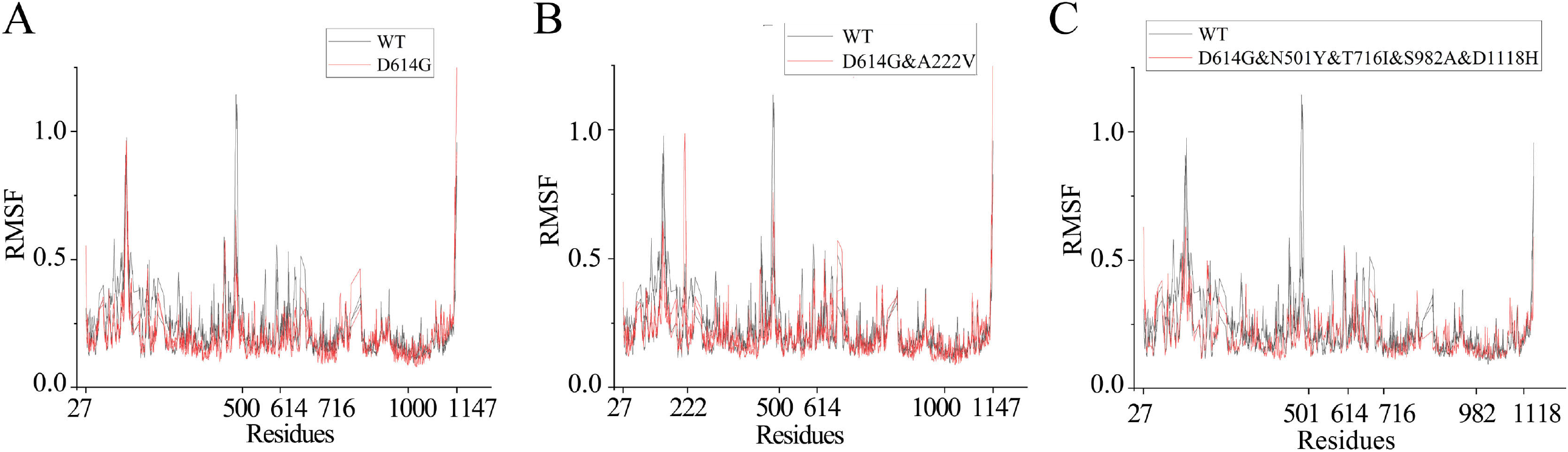
Flexibility evaluation of S protein and its mutants. A. Comparison of flexibility between WT and D614G. B. Comparison of flexibility between WT and D614G&A222V. C. Comparison of flexibility between WT and D614G&N501Y&T716I&S982A&D1118H.

Analysis of solvent accessible surface area (SASA) as a surface accessible to proteins provides information about the ability of proteins to interact with other molecules (Table S5). The whole molecular SASA of the WT, D614G, D614G&A222, and penta-site mutants was analyzed (Figure S1). The results showed that under the stable state, the SASA of the penta-site mutant was significantly higher than that of D614G and slightly higher than that of D614G&A222V and WT. The SASA values of D614G&A222V and WT were very close and fluctuated within a small range.

### No holistic change in the antigenicity of S protein and their mutants

The development of a vaccine is one of the most important means of long-term prevention and control of the virus. Antigenicity is an indicator of the ability of a peptide to bind to its complementary antibodies ^26^. The antigenicity was analyzed by Ellipro ^27^, a structure-based web tool that implements Thornton’s method and, together with a residue clustering algorithm, the MODELLER program and the Jmol viewer. The antigenicity of S protein was analyzed (Figure S2, Table S6), which showed that the antigenicity of S protein fluctuated highly in different local regions, but there was no significant change in the overall trend. This indicated that these mutations did not change the ability of the S protein to react with the antibodies. However, the potential for reinfection caused by the mutation remains a concern.

## Discussion

S protein is located on the surface of SARS-CoV-2, and it has been indicated that the S protein of SARS-CoV-2 has a characteristic Furin cleavage site at the boundary between S1/S2 subunits. ^2,5,28^ S protein mediates the entry of SARS-CoV-2 into cells through ACE2. The amino acid variation in the SARS-CoV-2 region will lead to the change of virus infection characteristics, which is related to the effective transmission of SARS-CoV-2 in the human population.

Mutation is one of the main mechanisms by which viruses are constantly changing due to genetic selection. Although most point mutations are neutral and do not alter the protein which the gene codes for, a few favorable mutations can give an evolutionary advantage to viruses ^29^. The D614G substitution in the gene encoding the spike protein emerged in late January or early February 2020 and emerged in Europe around 2020 February 22 ^17^. Since then, the D614G variant quickly became popular in Europe and by June 2020 became the dominant form of the virus circulating globally ^15,17,30^. The D614G mutation has been reported to change the S protein conformation and enhance the efficiency of protease cleavage at the S1/S2 subunit junctions, which may be one of the reasons for promoting its infection to the host and improving the transmission efficiency of SARS-CoV-2 ^31^. Studies on the D614G mutant of S protein at the atomic simulation level indicate that D614G has stronger adaptability and a higher transmissible carbon skeleton ^32^. A222V has recently been reported as a rapidly increasing genotype ^33,34^. Mutants that involve the penta-site mutants were first reported in the United Kingdom from October to November 2020, including two lineages of co-prevalence. The S protein mutants were single mutation of N501Y and multi-site mutation including N501Y, T716I, S982A, and D1118H. The two lineages spread rapidly after they emerged ^9^. 501Y Variant 2 was also named B.1.1.7 by COVID-19 Genomics Consortium UK (CoG-UK) ^35^. It should be pointed out that the S protein of B.1.1.7 also had other mutations including P681H and Y144 deletion in addition to the penta-site mutation. The mutation sites selected in this study were from the AutoVem tool. With the exception of P681H, which has not been structurally resolved in the protein used in this study, other mutations are not significant in the real-time monitoring of AutoVEM at present. This suggests that the adequacy and necessity of mutant phenotypes in the process of epidemiological transmission may be a question worth considering.

The changes in protein morphology caused by these mutations may provide a possible explanation for their roles in transmission (Figure 4, Figure S3). We analyzed the local structural changes in residues which are close to the mutation sites (no more than 10-nm distance). For D614G, it is located between the S1 and S2 subunits of the S protein. Two subunits of the S protein are cleaved by host cell proteases when the virus enters the host cell. Morphologically, the transition from aspartic acid to glycine eliminates a bulge on the protein surface (Figure 4A, Figure S3A), which may reduce steric hindrance during protease action and facilitate virus entry into cells. For the mutation site A222V, as the amino acid at the edge of the peptide chain, the change from alanine to valine extends the side chain of the amino acid, and both of them are hydrophobic. The positive increase of the hydrophobic effect of this change may contribute to the binding between the peptide chains (Figure 4B, Figure S3B). N501Y as the S protein RBD locus mutation has been closely watched, asparagine to tyrosine in the form not only produced a raised, and the side chain into a with active hydroxy benzene ring, change together with the presence of hydroxyl in the form of greatly improved its ability to react with the outside world (Figure 4C, Figure S3C), which could enhance the S protein of ACE2 ability. Similar to N501Y, T716I also increases a bulge on the surface of the protein. Meanwhile, the disappearance of the side chain hydroxyl group and the transformation into a hydrophobic side chain change the tendency of the amino acid to interact with the outside world from the formation of hydrogen bonds to hydrophobicity, which may have a positive effect on the presence of S protein on the surface of the viral membrane (Figure 4D, Figure S3D). S982A is located at the edge of the peptide chain, and like T716I, it loses the hydroxyl group of the side chain, and the side chain becomes hydrophobic, which may enhance the hydrophobicity between the peptide chains and facilitate the binding between the peptide chains (Figure 4E, Figure S3E). D1118H is located at the bottom of the S protein and mutates to form a large bulge. The side chain changes from carboxyl group to imidazole group, which may be affected by the binding between the peptide chains (Figure 4F, Figure S3F).

**Figure 4.**
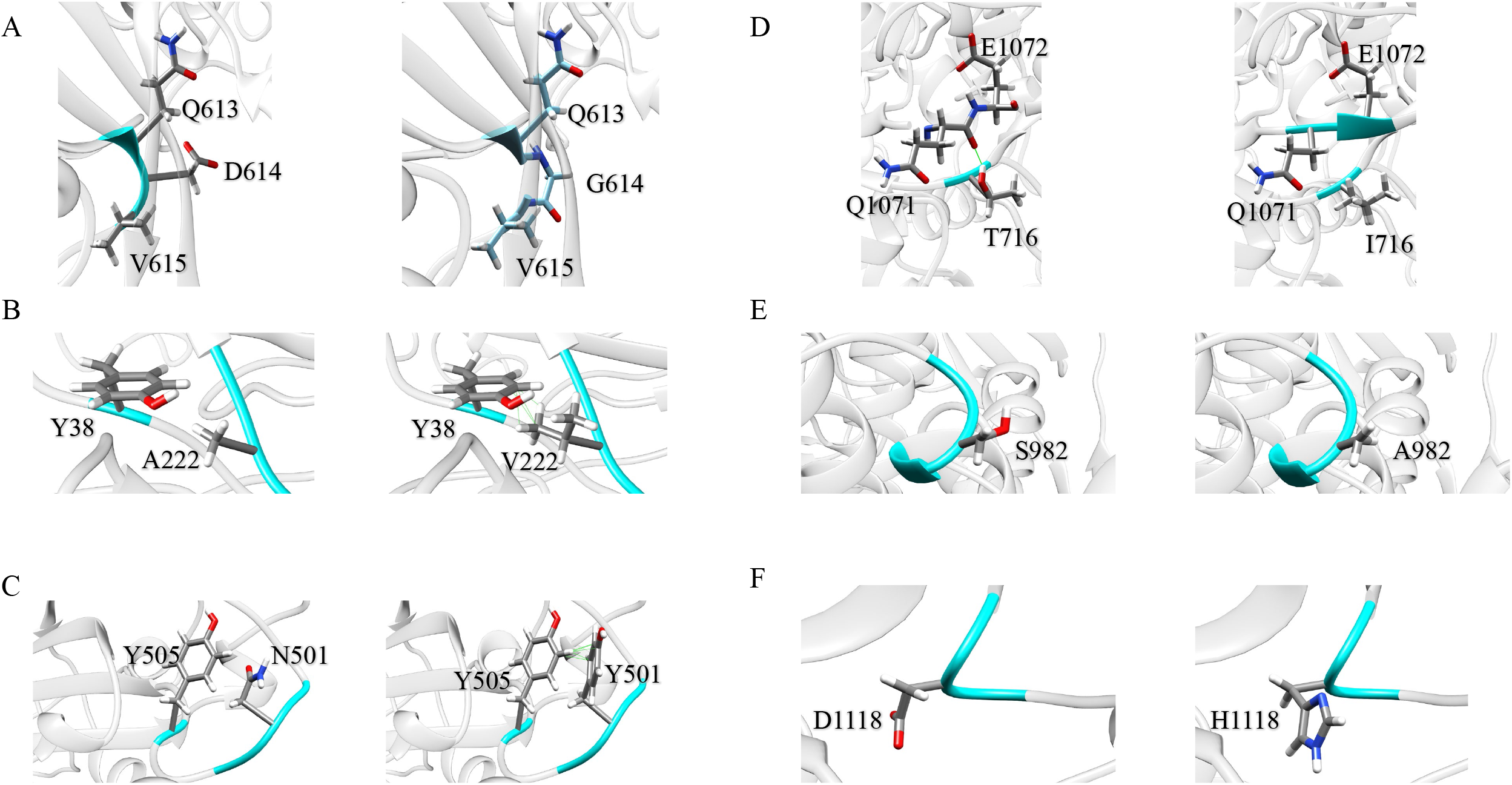
Local structural comparison between mutants and WT. The amino acid structure of 10 Å around the mutation site of the A chain of the mutant and WT is shown. The region where the mutation site is located is rendered as cornflower blue, the non-A chain region is rendered as cyan, and the other parts are rendered as pink. A. The amino acid mutation at position 614 showed significant morphological changes from D to G. The G614 side chain was shorter, which reduced the steric hindrance in this region, and the hole on the protein surface (Figure S3) could be observed. Meanwhile, the chemical properties of the side chains of D and G are also highly changed. The side chain of D has two highly active hydroxyl groups, while the side chain of G is inactive methyl group. B. The amino acid at position 222 mutated from A to V, increasing the volume of the amino acid at this position, and at the same time, it had interactions (van der Waals force, etc) with 38Y that were not existed before. Due to the similar atomic composition of side chains between A and V, these interactions should be caused by the reduction of residue distance. C. The mutation of N to Y at 501 causes a change in the size and morphology of the amino acid at this position. The amino acid extends outward, about 10 Å away from the 439N of the B chain, and is more likely to react with the protein receptor ACE2 above the virus. In addition, N501Y also exhibits multiple non-existent interactions with amino acids at position 506. D. The change of amino acid T to I at position 716 not only results in the loss of an active hydroxyl group, but also the loss of an internal protein interaction. The change of hydroxyl group to methyl group also leads to the change of hydrophilicity and hydrophobicity locally. E. The change of amino acid from S to A at the position 982 lost a hydroxyl group, resulting in the change of hydrophilic and hydrophobic properties and chemical properties. F. The change of amino acid D to H at 1118 did not lead to the change of internal protein interaction, but the change of side chain affected the chemical properties of this region. At the same time, the ring structure of histidine side chain also changes the steric hindrance in this region.

During the analysis phase in this study, RMSD values were observed to be in a stationary state, indicating that the structure fluctuated around a stable mean conformation. Therefore, it is meaningful to evaluate the fluctuation of local structure^25,36^. During the stable period (the latter 50ns) of the simulation process, the two proteins and their mutants showed different properties. For S protein, the RMSD (based on the analysis of Cα) of D614G, D614G&A222V, and penta-site mutant has a certain decline compared with wild type, this suggests that the D614G mutation promoted S protein stability ^37^, the promotion of the stability is proportional to the decline degree ^18^. This is also consistent with current epidemiological data. Besides, the RMSD of all D614G carrying mutants decreased, the stability was increased. And the addition of A222V and N501Y&T716I&S982A&D1118H respectively increases the values of SASA and R_g_. It suggested possibly that the D614G single mutation resulted in a decrease in RMSD, while A222V&D614G and penta-site showed an increase in SASA and R_g_. The results probably suggested that D614G improved the stability of S protein, while A222V and N501Y&T716I&S982A&D1118H both improved the ability of protein to react with the outside environment.

The average number of hydrogen bonds of the mutant D614G and D614G&A222V in the stable state was higher than that of the WT and the penta-site mutant. The number of hydrogen bonds could reflect the stability of the protein to some extent. It should be noted that interaction does not involve hydrogen bond formation alone. Although the number of hydrogen bonds in the D614G&A222V mutant was higher than that in the D614G mutant and the penta-site mutant, the RMSD was lower. This suggests that the stability of S protein is not only influenced by hydrogen bonding, but other interactions such as hydrophobicity may also play important roles in stability. The analysis of SASA (habitual accessible surface areas) provides evidence to explain this phenomenon. Compared with D614G, D614G&A222V, and the penta-site mutant has a slightly higher value of SASA, which indicates that D614G&A222V and the penta-site mutant have higher protein interaction ability ^38^ while retaining the stability brought by the mutation at 614, and is more likely to react with the outside environment. This further enhanced the efficiency of transmission, which is consistent with epidemiological data.

Our study provides a mechanistic hypothesis for the change in transmissibility caused by mutations in the SARS-CoV-2 genotype. Although these mutations and their possible consequences of re-infection are disturbing ^39^, fortunately, our analysis of the antigenicity of S proteins suggests that these mutations do not significantly alter their ability to react with antibodies. However, a recent study has revealed the potential pathogenicity of the S protein itself, suggesting that mutations in the S protein itself may have a more direct effect on COVID-19 than commonly believed^40^. The prevention and control of SARS-CoV-2 is a long process. With the emergence of more and more mutants, the mechanism of “genotype -- phenotype -- epidemiological characteristics” caused by mutations will become one of the key directions of future research. It is worth emphasizing that, based on the real-time monitoring of the mutant strains of the pandemic virus, the S protein linkage mutation was studied as a whole rather than a single mutation site, which was more in line with the actual transmission of the virus and provided some reference for the continuous exploration of molecular dynamics for the deeper understanding of SARS-CoV-2.

## Resource Availability

### Lead Contact

Further information and requests for resources and reagents should be directed to and will be fulfilled by the Lead Contact, Hongli Du

### Materials Availability

This study did not generate new unique reagents.

### Data and Code Availability

Raw data tables for RMSD, RMSF, RG, SASA, H-Bond, and Antigenicity are provided in the supplementary information. The simulation data is stored in the HPC hardware housed at the Tianhe-2 and can be shared with the corresponding author on request.

## Methods

The cryo-electron microscopic structures of the wide type S protein (PDB ID: 6×6P) and ORF3a (PDB ID: 7KJR) were obtained from the Protein Data Bank ^41^. The key mutation information of S protein was previously studied by Xi et al 2021. The sequence information comes from the A chain of the protein. The point mutation of protein was conducted by MOE2019.

Using the structures obtained from PDB as initial templates, we induced mutations in Molecular Operating Environment 2019 to obtain D614G, D614G&A222V and the penta-site mutants. For each system, the MD simulations were performed using Gromacs 2018.3, and the Gromacs 54A7 force field was selected ^42^. All structures were solvated with SPCE ^43,44^ water molecules in a cubic simulation box, and Na^+^ and Cl^-^ ions were added to neutralize the system. We first performed a 5000-step energy minimization using the steepest descent method, followed by three processes in the MD simulation: NVT, NPT, and production. The NVT ensemble (constant particle number, volume, and temperature) in 300 K for 100 ps, followed by 100 ps in the NPT ensemble (constant particle number, pressure, and temperature) at 1 atm. The above simulations are all locational constraints, in which all the bonds are confined to the protein to relax the water around the protein and reduce the entropy of the system. Select Parrinello□Rahman as the barometric regulator and V-Resale as the thermostat.

Finally, a 100-ns production run was performed at 300 K. LINCS algorithm was used to constrain all the bonds, and the cutoff distances of 12 Å for the long-range electrostatic through the Particle Mesh Ewald (PME) method. A time step of 0.002 ps was selected in the simulation without constant force being applied.

The antigenicity information is analyzed using the Ellipro ^27^ server with the default parameters.

The biological macromolecules presented in this paper were drawn with PyMol V2.3.2 and UCSF Chimera 1.15^45^, and the statistical charts were drawn with Origin 2018.

## Acknowledgments

This work was supported by the National Key R&D Program of China (2018YFC0910201), the Key R&D Program of Guangdong Province (2019B020226001), and the Science and the Technology Planning Project of Guangzhou (201704020176).

## Author contributions

J.R.L. designed the experiments and analyzed the data. B.X. and Z.C. did the calculation of simulation. B.Z. did the calculation of antigenicity and helped finish the manuscript. Y.Z. helped finish the manuscript. R.Q. and S.Z. helped helped with the introduction and disposal data. H.D. and P.G. designed the project and helped finish the manuscript.

## Supplemental Information

Table S1 RMSD OF S

Table S2 H-bond OF S

Table S3 Rg OF S

Table S4 RMSF OF S

Table S6 SASA OF S

Table S6 Antigenicity OF S

Figure S1 The SASA of S protein and mutants Figure S2 The Antigenicity of S

Figure S3 Protein surface and residues of mutants

A. D614G; B. T716I; C. A222V; D. N501Y; E. S982A; F. D1118H

