## Supplementary figures and images for "Molecular dynamics simulation study reveals effects of key mutations on spike protein structure in SARS-CoV-2"

### Supplemental Figures

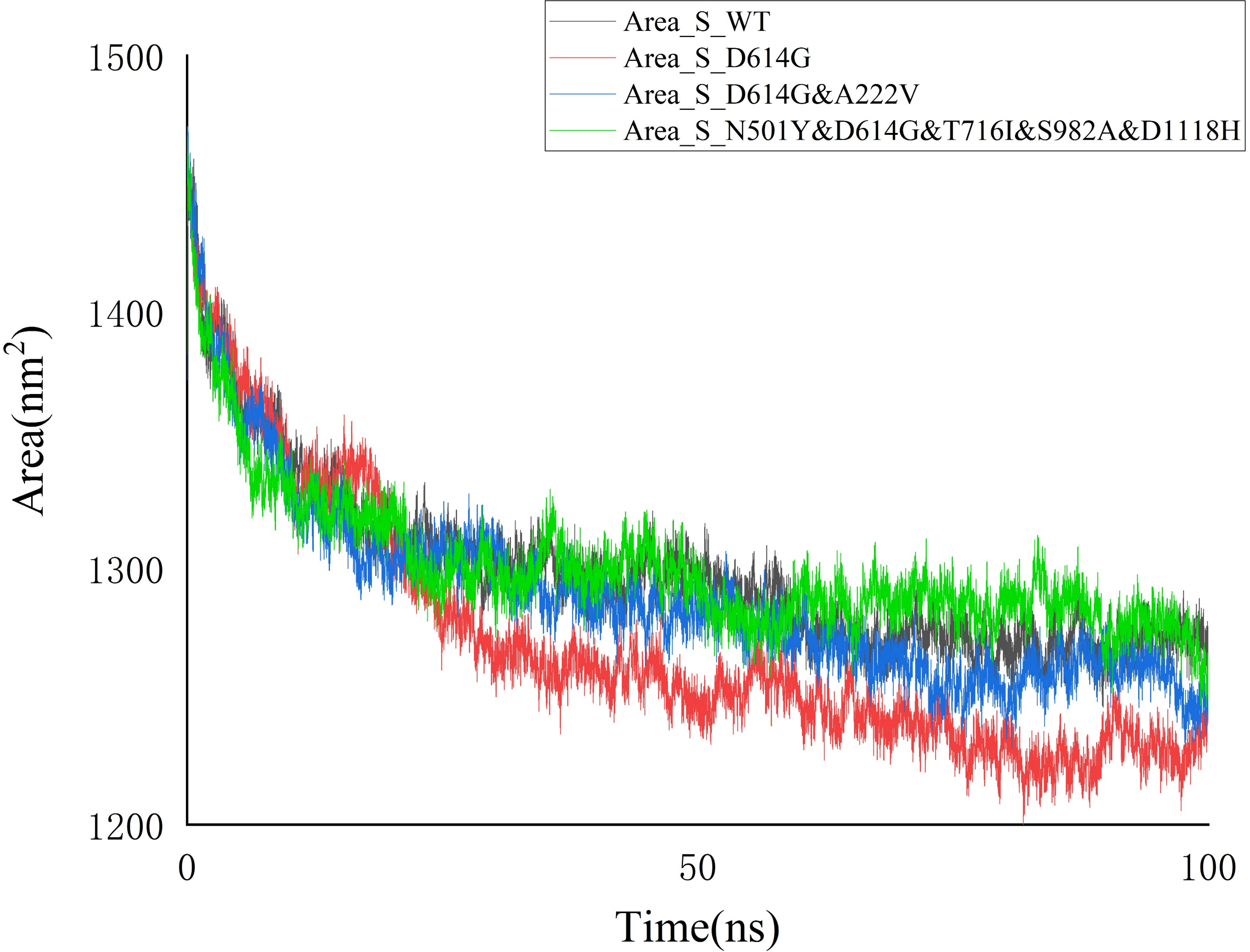

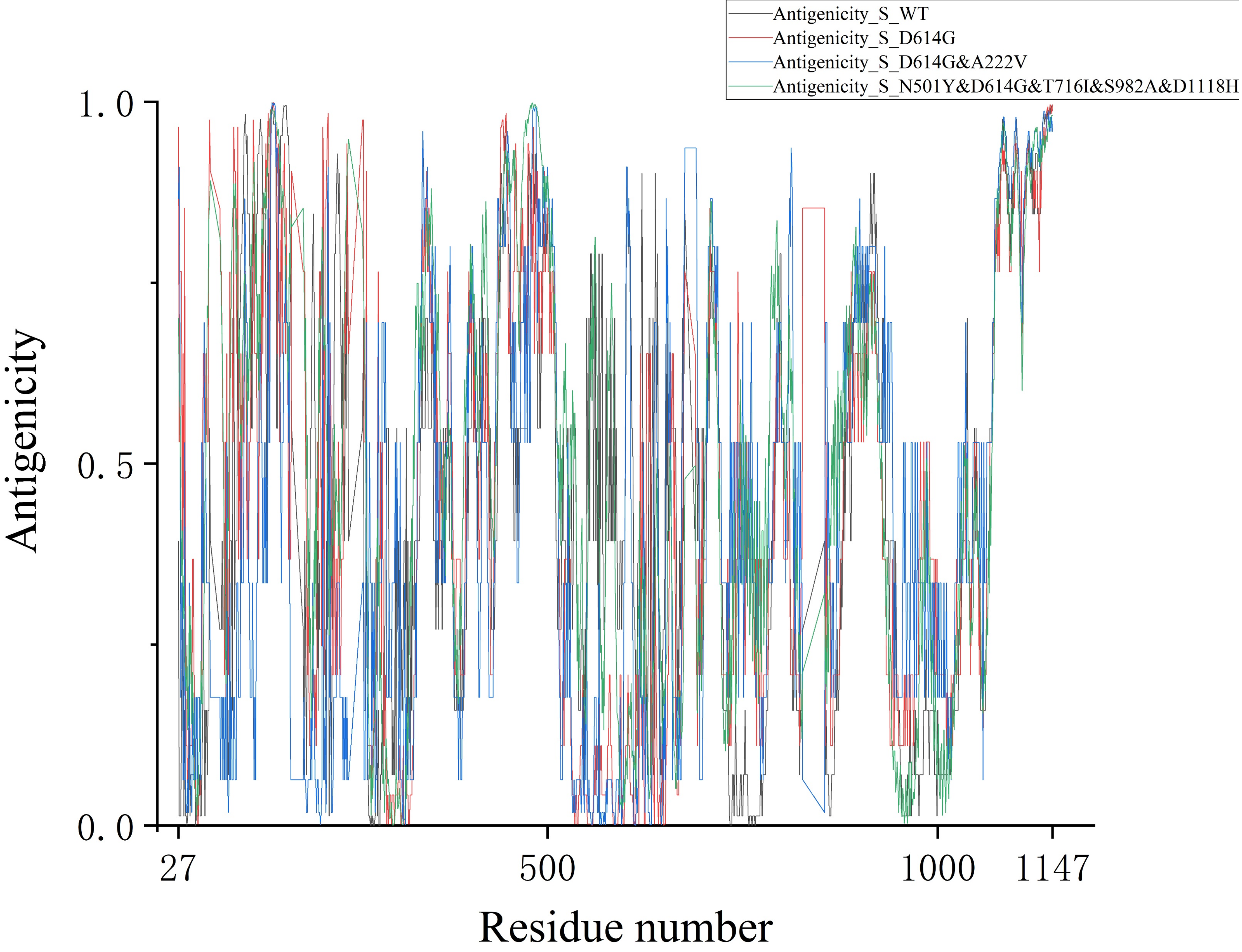

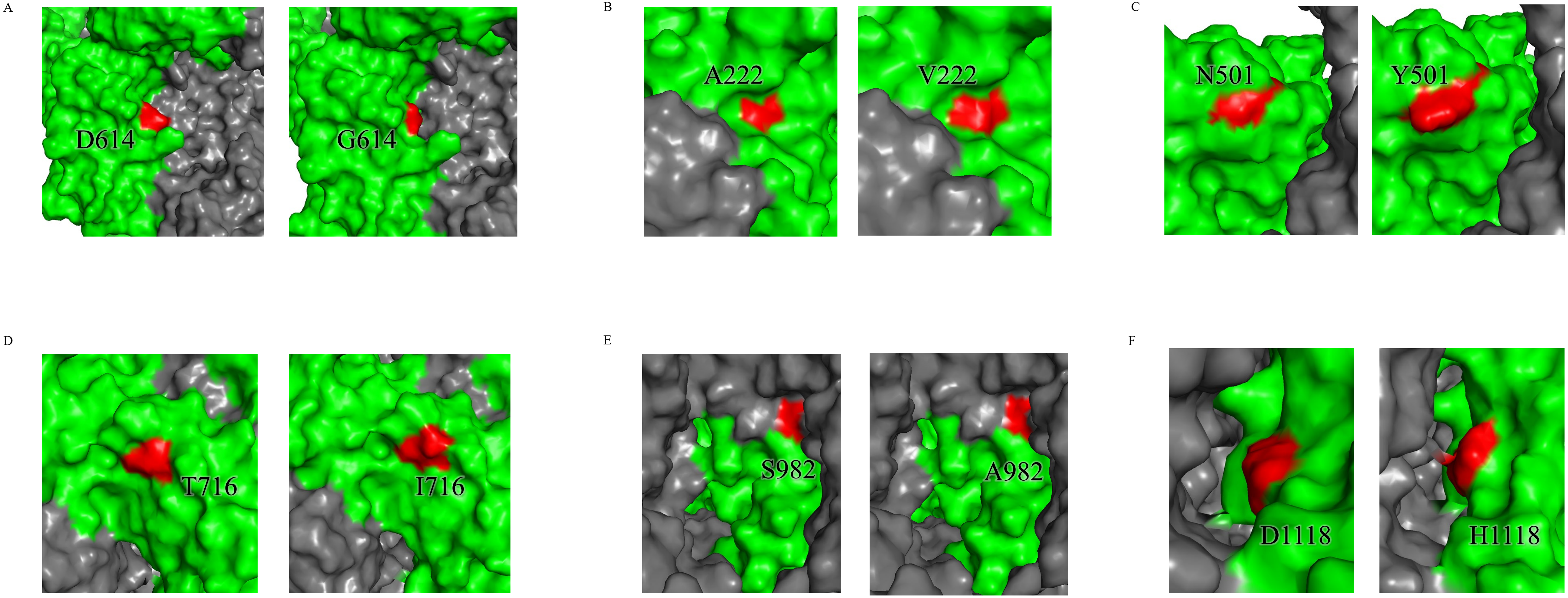
